# Identification of senescence-related hub genes and potential therapeutic agents for chronic heart failure

**DOI:** 10.1101/2025.06.21.660877

**Authors:** Hao Xie, Zuyuan You, Bin Liao, Juyi Wan, Mingbin Deng, Qi Yang

**Affiliations:** Department of Cardiovascular Surgery, The Affiliated Hospital, Southwest Medical University, Metabolic Vascular Diseases Key Laboratory of Sichuan Province, Key Laboratory of cardiovascular remodeling and dysfunction, Luzhou 646000,Sichuan, P. R. China; Key Laboratory of Medical Electrophysiology, Ministry of Education & Medical Electrophysiological Key Laboratory of Sichuan Province, (Collaborative Innovation Center for Prevention of Cardiovascular Diseases), Institute of Cardiovascular Research, Southwest Medical University, Luzhou 646000, Sichuan, P. R. China; Department of Cardiovascular Surgery, Chongqing University Three Gorges Hospital, Chongqing 404000.China

**Author notes:** **Corresponding Authors:** Mingbin Deng and Qi Yang, (MD); (QY). (HX); (ZY), (BL); (JW). Hao Xie and Zuyuan You have contributed equally to this work. Mingbin Deng and Qi Yang are co-corresponding authors.

**Keywords:** Chronic heart failure, Senescence, Diagnostic marker, Molecular docking

## Abstract

**Background:** Chronic heart failure (CHF), the terminal phase of cardiovascular disease progression, has emerged as an increasingly severe global public health concern. Despite current therapeutic approaches aimed at symptom relief, their long-term effectiveness remains limited, urgently necessitating the exploration of novel treatment strategies. This study endeavored to explore the role of cellular senescence in CHF, identify the characteristic genes linked to cellular senescence, and predict potential therapeutic agents.

**Methods:** We acquired CHF-related datasets from the Gene Expression Omnibus database and cell senescence-related genes from the CellAge database to identify differentially expressed cell senescence-related genes. We then conducted Gene Ontology and Kyoto Encyclopedia of Genes and Genomes analyses to elucidate the functions of these differentially expressed genes, constructed a protein–protein interaction network, and screened hub genes. Using receiver operating curve (ROC) analysis, we developed a diagnostic model based on these hub genes. Furthermore, we constructed networks for the hub genes involving miRNAs, lncRNAs, transcription factors, and drugs. Subsequently, we explored the potential mechanism of action of metformin in the treatment of CHF through molecular docking studies. Lastly, we verified the expression of the hub genes in a doxorubicin-induced CHF model in rats. **Results** We ultimately identified nine hub genes associated with cellular senescence: *STAT1*, *MMP9*, *MAP2K1, SOCS1*, *SDC1*, *MET*, *EIF4EBP1*, *ATF3*, and *NAMPT*. These genes exhibited significant differential expression between CHF and normal tissues. The constructed diagnostic model demonstrated robust diagnostic performance in ROC curve analysis, with an area under the curve exceeding 0.7, thereby providing biomarker support for early CHF diagnosis. Furthermore, we identified a regulatory network comprising 94 lncRNAs, 63 miRNAs, and 11 transcription factors and screened 42 potential therapeutic drugs. Subsequent molecular docking simulations revealed that metformin could effectively bind to the hub genes. Finally, we validated the expression of some hub genes in a rodent CHF model, in which gene expression regulation varied across diverse experimental settings.

**Conclusions:** Our research identified nine genes cellular senescence-related genes with potential roles in CHF pathogenesis, offering fresh perspectives concerning the diagnosis and treatment of CHF.

## Background

Chronic heart failure (CHF) describes a complex clinical syndrome resulting from abnormal alterations in cardiac structure or function with diverse causes, ultimately impairing the heart’s contractile and diastolic functions[1, 2]. The primary manifestations of CHF include dyspnea, fatigue, and fluid retention (manifesting as pulmonary congestion, systemic congestion, and peripheral edema)[3]. A recent study revealed that, as of 2022, more than 64 million people globally have heart failure[4]. Given its exceptionally high incidence and mortality rates, CHF represents a challenging and difficult-to-manage cardiovascular disease[5].

The pathogenesis of CHF is notably intricate and multidimensional, generally encompassing myocardial remodeling, abnormal myocardial energy metabolism, and neuroendocrine system hyperactivity[6]. Currently, CHF treatment primarily aims to alleviate symptoms by reducing the cardiac load and modulating the neuroendocrine system[7, 8]. However, the long-term efficacy of this strategy is suboptimal, posing certain clinical limitations. Consequently, in-depth research is needed to elucidate the mechanisms underlying CHF and identify potential drug targets for this condition.

Cellular senescence, as a self-defense mechanism in the natural evolutionary process of cells, has been extensively demonstrated to participate in pathophysiological processes such as injury and cancer[9, 10]. Previous studies revealed that senescent cells can accumulate and induce more severe senescence through the autocrine or paracrine secretion of senescence-associated phenotypes, ultimately leading to the functional impairment of organs such as the heart[11]. The proportion of senescent cardiomyocytes continues to increase with age. Senescent cardiomyocytes represent the biological basis for the development of heart failure[12]. According to research, Rb1 and Meis homeobox 2, as cell cycle inhibitors, can accelerate the aging of cardiomyocytes. However, by inhibiting these factors, the proliferative capacity of cardiomyocytes in adult rats was significantly enhanced, the infarct size after myocardial infarction was reduced, and the massive loss of cardiomyocytes and heart failure was alleviated[13]. These findings offer valuable insights into the mechanisms of myocardial cell senescence and CHF repair while also revealing potential therapeutic approaches.

Research in a senescence mouse model indicated that the induction of endothelial cell senescence through high-fat/high-salt diet consumption triggers endothelial cell inflammation, leading to the typical symptoms of CHF in mice, including left atrial dilation, left ventricular hypertrophy, and fibrosis. This suggests a relationship between endothelial cell senescence and CHF[14]. Different immune cells in human heart tissue directly regulate inflammatory responses and cardiomyocyte senescence, playing varying roles in CHF[15]. Studies revealed that senescent monocytes/macrophages produce various proinflammatory factors, and macrophages can release the inflammatory factors IL-4 and IL-3 by promoting the phosphorylation of signal transducer and activator of transcription 3 (STAT3), leading to cardiomyocyte hypertrophy. Persistent myocardial hypertrophy can result in maladaptive ventricular remodeling, which is considered the main cause of CHF[16]. Senescent CD4^+^ T cells secrete a large amount of interferon-γ, which infiltrates the heart and causes myocardial inflammation and stress response, potentially eroding cardiac arterial plaques[17]. Research found that senescent T cells in patients with coronary heart disease can trigger a decline in left ventricular function and accelerate disease progression. This suggests that senescent immune cells increase the risk factors for CHF via multiple pathways[18]. Resveratrol exerts anticellular senescence effects by regulating the Akt/Bad/Bcl-2 pathway, thereby ameliorating the pathological changes of CHF[19]. Mice with IL-7 knockout exhibited significantly improved cardiac function and reduced cardiac hypertrophy, identifying IL-7 as a promising therapeutic target for CHF treatment[20]. Thus, the pharmacological clearance of senescent cells might represent a potential treatment strategy for CHF. However, senescence is a complex, multifactorial physiological process, highlighting the need for future research to develop CHF treatments based on anticellular senescence strategies and explore new prevention and treatment strategies for CHF.

In this background, the present study combined transcriptomics analysis and animal modeling to identify genes associated with cellular senescence with relevance to CHF and explore potential therapeutic agents. Through systematic analysis of public gene expression databases, such as Gene Expression Omnibus (GEO) and CellAge, we identified differentially expressed genes (DEGs) and elucidated their biological significance in CHF via functional enrichment analysis and constructed protein–protein interaction (PPI) networks. This multifaceted research methodology could facilitate a deeper comprehension of the pathogenesis of CHF and offer a theoretical foundation for future clinical treatments.

## Methods

### Data collection and processing

The series matrix files of GSE84796 and GSE76701 were downloaded from the GEO database (http://www.ncbi.nlm.nih.gov/geo)[21]. The expression profile data for GSE84796 comprise ten CHF heart and seven healthy patient heart tissues. The expression profile data for GSE76701 consist of four CHF heart and four healthy patient heart tissues. In total, 886 cell senescence-related genes were downloaded from the CellAge database (https://genomics.senescence.info/cells/). The analysis flowchart for this study is presented in Fig. 1.

**Fig. 1.**
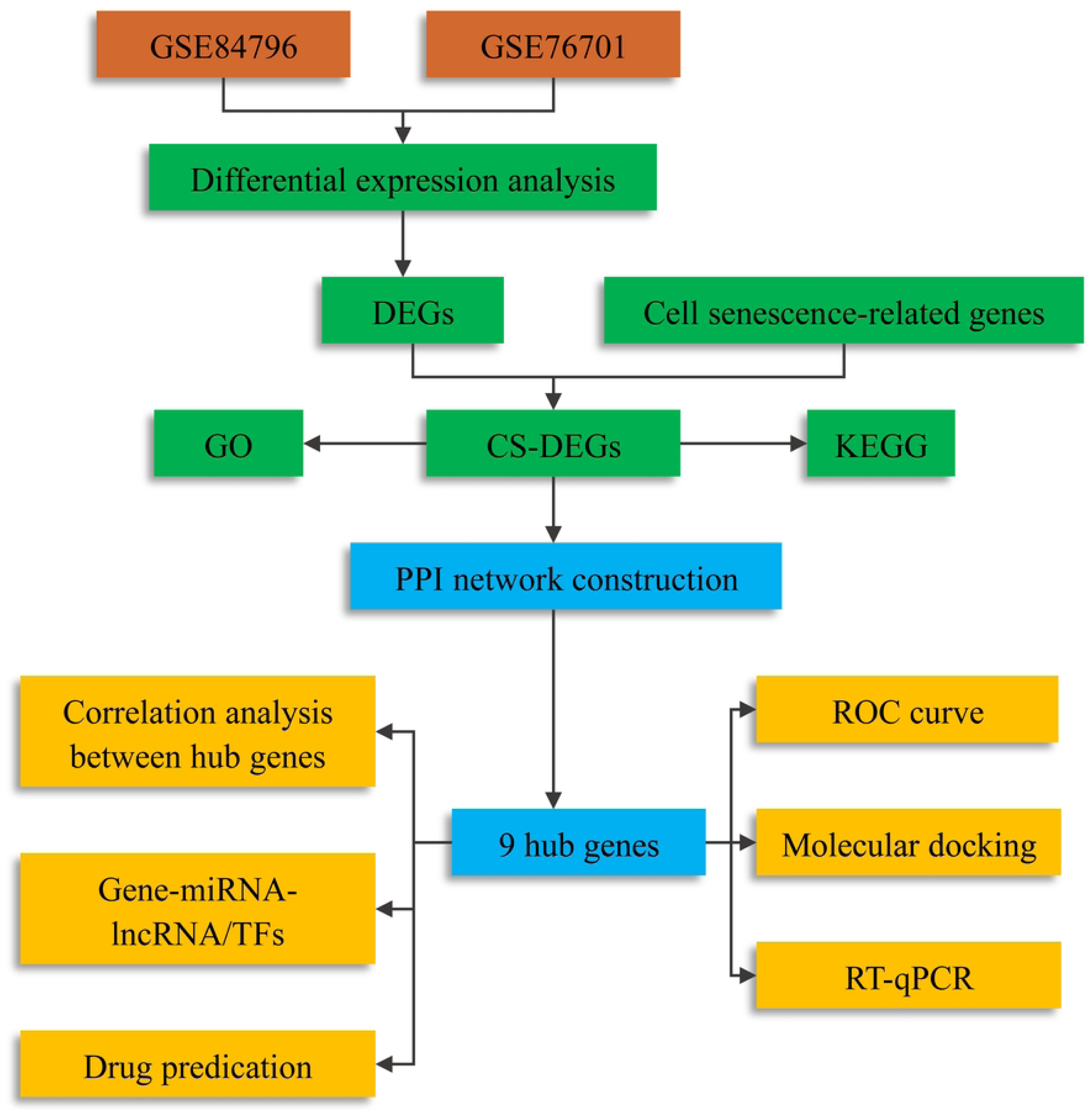
Overall study process.

### Identification of DEGs

The raw data were processed using R software (The R Foundation for Statistical Computing, Vienna, Austria). The “limma” package was employed for differential expression analysis using adjusted p-value < 0.05 and |log_2_FC| > 0.585 as the screening criteria. The “ggplot2” and “pheatmap” packages were used to visualize the expression information of the DEGs as volcano plots and cluster heatmaps[22, 23]. Subsequently, we utilized Venn diagrams to intersect the filtered DEGs with the cell senescence-related gene set, to identify common DEGs associated with both CHF and cell senescence.

### Functional enrichment analysis

To identify the biological functions of common DEGs, we conducted Gene Ontology (GO) analysis and Kyoto Encyclopedia of Genes and Genomes (KEGG) enrichment analysis. In this study, we imported common DEGs into the DAVID database (https://david.ncifcrf.gov/). We then performed GO and KEGG analyses across three modules: biological process (BP), cellular component (CC), and molecular function (MF). Genes were considered significantly enriched at P < 0.05. Subsequently, the enrichment analysis data were visualized using Metascape (https://metascape.org/gp/index.html#/main/step1).

### Analysis of the PPI network and screening of hub genes

The STRING database (https://cn.string-db.org/) was utilized to identify interactions between known and predicted proteins. After importing common DEGs, the minimum required interaction score was set to 0.4, and the PPI network was drawn. Subsequently, the calculation results were imported into Cytoscape (version 3.9.1, https://cytoscape.org/)[24]. The CytoHubba plugin within Cytoscape software offers 12 importance ranking algorithms, namely the maximum neighborhood component, density of maximum neighborhood component, maximum cluster centrality, degree, edge percolation component, bottleneck, eccentricity, closeness, radiance, betweenness, stress, and repeat factor. These algorithms enable the assignment of importance scores to each node.

### Receiver operating characteristic (ROC) curve validation and correlation analysis

In the combined dataset of GSE84796 and GSE76701, box plots of gene expression profiles for hub genes were generated using the “ggplot2” package in R. To verify the accuracy of the filtered hub genes, we conducted ROC curve analysis and calculated area under the curve (AUC) using the “pROC” package. AUC > 0.7 represented an ideal diagnostic value, and it was used for further analysis. Subsequently, the correlation between the expression of each hub gene pair was calculated. Scatter plots were drawn for the gene pairs exhibiting the strongest positive and negative correlations.

### Prediction of the gene–miRNA–lncRNA and gene–transcription factor (TFs) regulatory networks

We utilized the NetworkAnalyst database (https://www.networkanalyst.ca/) to predict the upstream regulatory miRNAs and lncRNAs of the hub genes. Subsequently, we predicted the TFs associated with the hub genes. Cytoscape was employed to construct the mRNA–miRNA–lncRNA and mRNA–TF interaction networks.

### Drug prediction for hub genes

By searching the Drug SIGnatures DataBase (DSigDB, https://dsigdb.tanlab.org/DSigDBv1.0/), candidate drugs targeting hub genes were screened, and Cytoscape was utilized for visualization[25].

### Molecular docking

First, the protein structures of the hub targets were downloaded from the Protein Data Bank (https://www.rcsb.org/) using the following criteria: originating from *Homo sapiens*, lacking mutations, possessing higher resolution, and containing small-molecule ligands. Subsequently, the structures of the compounds were downloaded from PubChem (https://pubchem.ncbi.nlm.nih.gov/). Using AutoDockTools (v1.5.7, https://autodocksuite.scripps.edu/adt/), we performed hydrogenation, dehydration, and other preprocessing on the hub target proteins and compound ligands. Finally, molecular docking was conducted using AutoDockVina (https://vina.scripps.edu/). The results of the molecular docking were visualized using PyMOL 3.0.3 software (https://www.pymol.org/)[26].

### Construction of a rat model of CHF

This study was approved by the Animal Experimental Ethics Committee of Southwest Medical University. Four-week-old male Sprague–Dawley rats were obtained from the Animal Center of Southwest Medical University (Luzhou, China) and raised in a pathogen-free environment. Each rat in the CHF group received intraperitoneal injections of doxorubicin (HY-15142 A, 2.5 mg/kg/week, MedChemExpress, Monmouth Junction, NJ, USA) for 6 weeks[27]. The control rats received injections of an equal volume of sterile saline at the same time points.

### Echocardiography

Two-dimensional targeted M-mode tracing was performed at the papillary muscle level using an echocardiography system (iE33, Philips, Amsterdam, Netherlands) equipped with a 124-MHz transducer (Philips). A series of cardiac parameters were measured and calculated. For each measurement, the average value of three consecutive heartbeats was used[28].

### Histological and morphological analyses

After rats were sacrificed, their hearts were excised, perfused with cold PBS, and rinsed to remove blood on the surface. The moisture was absorbed with filter paper, and the hearts were then fixed in 10% formaldehyde for 24 h. Subsequently, the tissues were embedded in paraffin, sectioned (3 μm), and subjected to hematoxylin and eosin staining of the left ventricular tissue[29]. Images of the tissues were taken and analyzed using ImageJ (US National Institutes of Health, Bethesda, MD, USA).

### RT-qPCR

Total RNA was extracted from the left ventricular myocardial tissue of rats using TRIzol (Thermo Fisher Scientific, Waltham, MA, USA) and reverse-transcribed into cDNA using a reverse transcription kit (Roche, Basel, Switzerland). RT-qPCR was conducted using SYBR Green (Roche)[30]. The primers used for amplification are listed in Supplementary Material S1. The expression of the target genes relative to GAPDH gene expression was presented as 2^−ΔΔCt^.

### Statistical analysis

Statistical analysis was conducted using R software version 4.2.2. Wilcoxon’s signed rank test or Student’s *t*-test was employed to analyze the differences between the two groups. The Pearson or Spearman correlation test was conducted to determine the correlations between variables. Experimental data were expressed as the mean ± SD, and *P* < 0.05 denoted statistical significance. Bar charts were plotted using GraphPad Prism (version 6.0, GraphPad Software, Boston, MA, USA).

## Results

### Identification of cell senescence-related DEGs (CS-DEGs) in CHF

To investigate DEGs in CHF, we downloaded raw gene expression data from the GSE84796 and GSE76701 datasets in the GEO database. The basic information of these two datasets is listed in Table 1. Following data preprocessing and cleaning, we adjusted for batch effects (Fig. 2A, B) and normalized the expression matrices of both datasets. The trends in the box plots were almost straight lines (Fig. 2C, D).

**Fig. 2.**
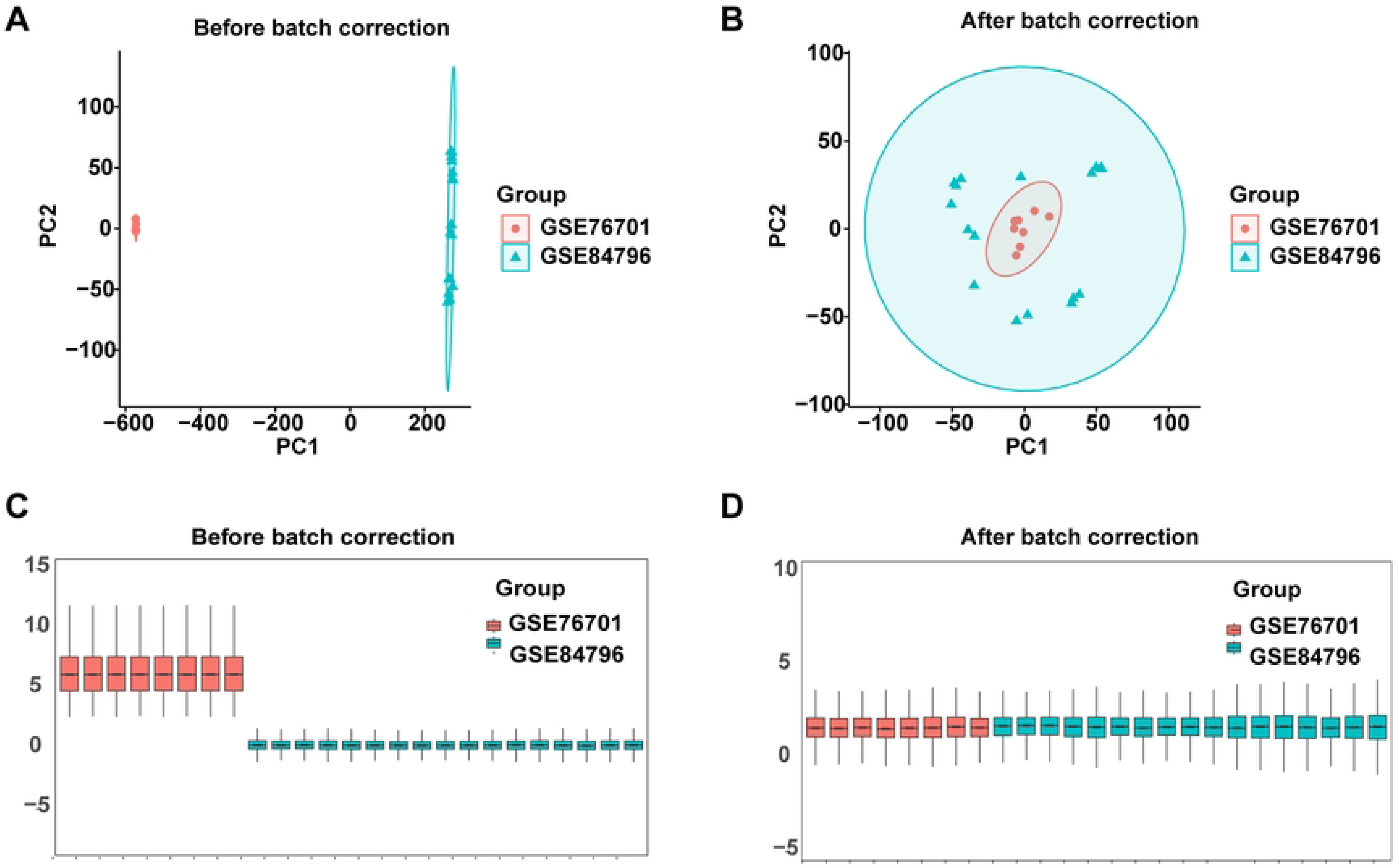
Principal component analysis (PCA) of the CHF and control groups using the combined microarray dataset. **A** PCA plot showing the expression of the combined microarray dataset (GSE84796 and GSE76701). **B** PCA plot showing the combined microarray dataset (GSE84796 and GSE76701) after removing batch effects. **C** Box plots before homogenization. **D** Box plots after homogenization.

Via differential analysis, 1223 DEGs were identified in the combined dataset of GSE84796 and GSE76701, of which 764 DEGs were upregulated and 459 DEGs were downregulated. The DEGs are displayed in the volcano plot in Fig. 3A, and the top 20 upregulated and downregulated DEGs are presented in the heatmap in Fig. 3B. To screen for cell senescence-related genes with differential expression in CHF, we intersected the DEGs in CHF with the 866 genes encoding cell senescence-related proteins listed in the database, resulting in the identification of 47 CS-DEGs (Fig. 3C).

**Fig. 3.**
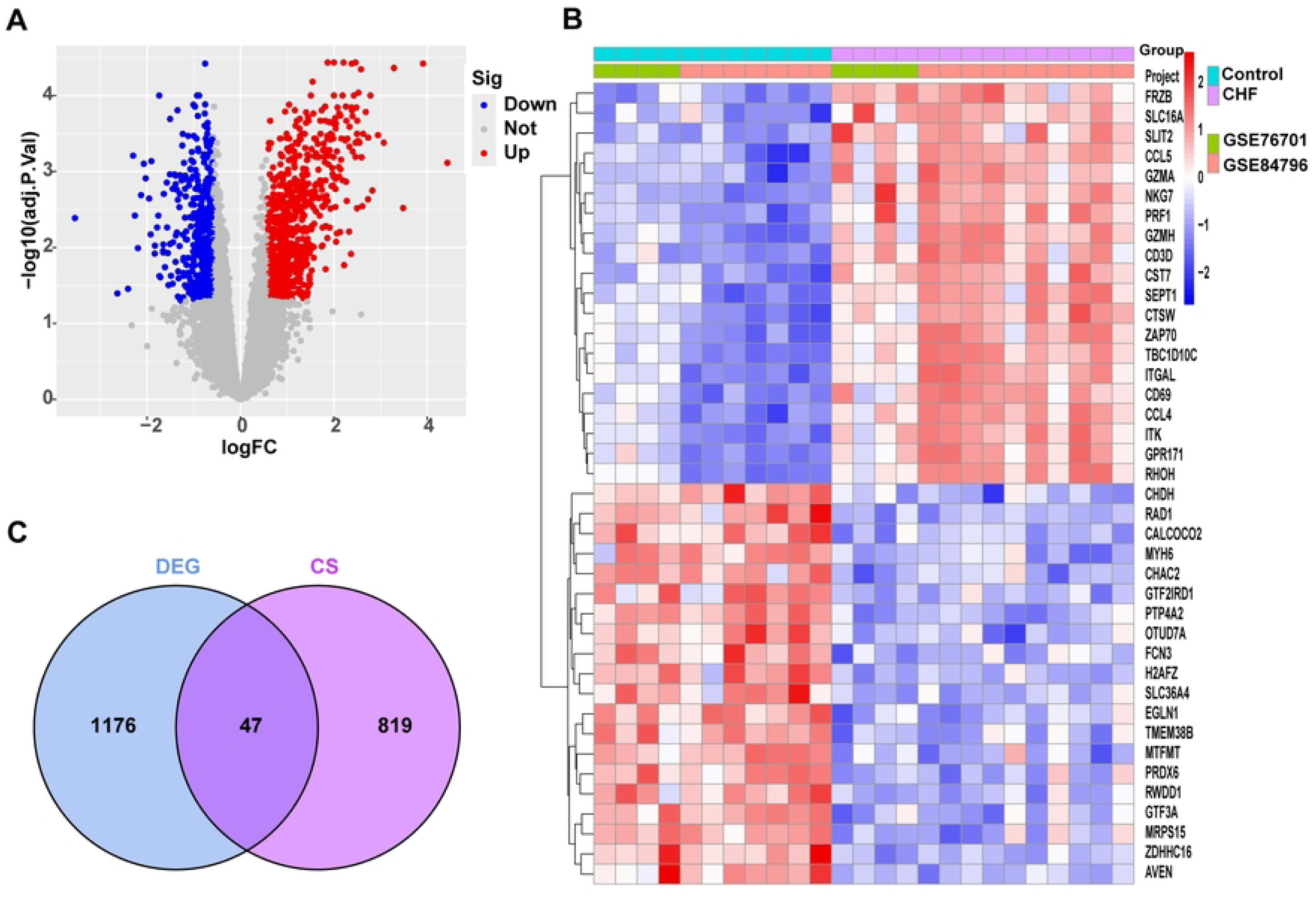
DEGs in the CHF and control groups in the combined microarray dataset (GSE84796 and GSE76701). **A** Volcano plot of DEGs in patients with CHF and matched controls. **B** Heatmap of the top 20 significantly upregulated or downregulated DEGs. **C** Venn diagram showing the intersection of DEGs with cell senescence-related (CS)-DEGs.

### Functional analysis of the CS-DEGs

Next, GO and KEGG enrichment analyses were conducted on 47 CS-DEGs to uncover their functions. The results revealed that CS-DEGs were primarily enriched in the positive regulation of the apoptosis process and protein phosphorylation (BP); the cytoplasm and mitotic spindle (CC); and activities related to protein binding, protein tyrosine kinase activity, and ubiquitin–protein ligase binding (MF, Fig. 4A). In the KEGG enrichment analysis, these 47 CS-DEGs were found to be enriched in diseases such as cancer, malaria, and hepatitis B and in pathways including the insulin, JAK– STAT, and Toll-like receptor signaling pathways (Fig. 4B).

**Fig. 4.**
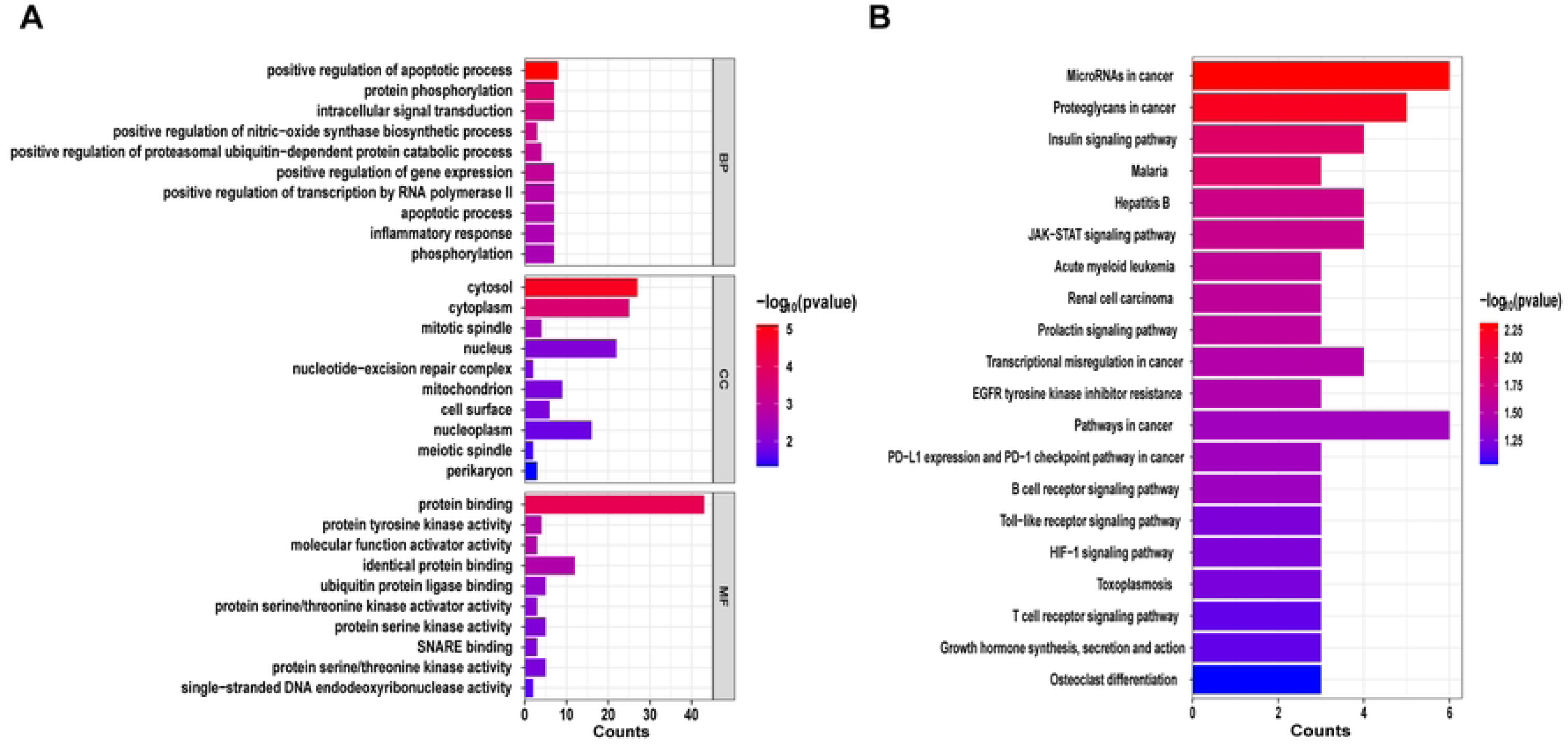
Functional enrichment analysis of CS-DEGs. **A** GO enrichment analysis of CS-DEGs. **B** KEGG pathway enrichment analysis of CS-DEGs.

### Construction of the PPI network and identification of hub genes

To further explore the potential relationships among the proteins encoded by CS-DEGs and identify the hub genes, a PPI network for CS-DEGs was constructed using the STRING database (Fig. 5A). Subsequently, the importance scores of each gene were calculated using 12 algorithms integrated within the CytoHubba plugin. For each algorithm, the top 20 genes were selected, and the intersection of these top 20 genes across all algorithms was determined, ultimately identifying nine hub genes: *STAT1*, *MMP9*, *MAP2K1*, *SOCS1*, *SDC1*, *MET*, *EIF4EBP1*, *ATF3*, and *NAMPT* (Fig. 5B). Detailed information on the hub genes is presented in Supplementary Material S2.

**Fig. 5.**
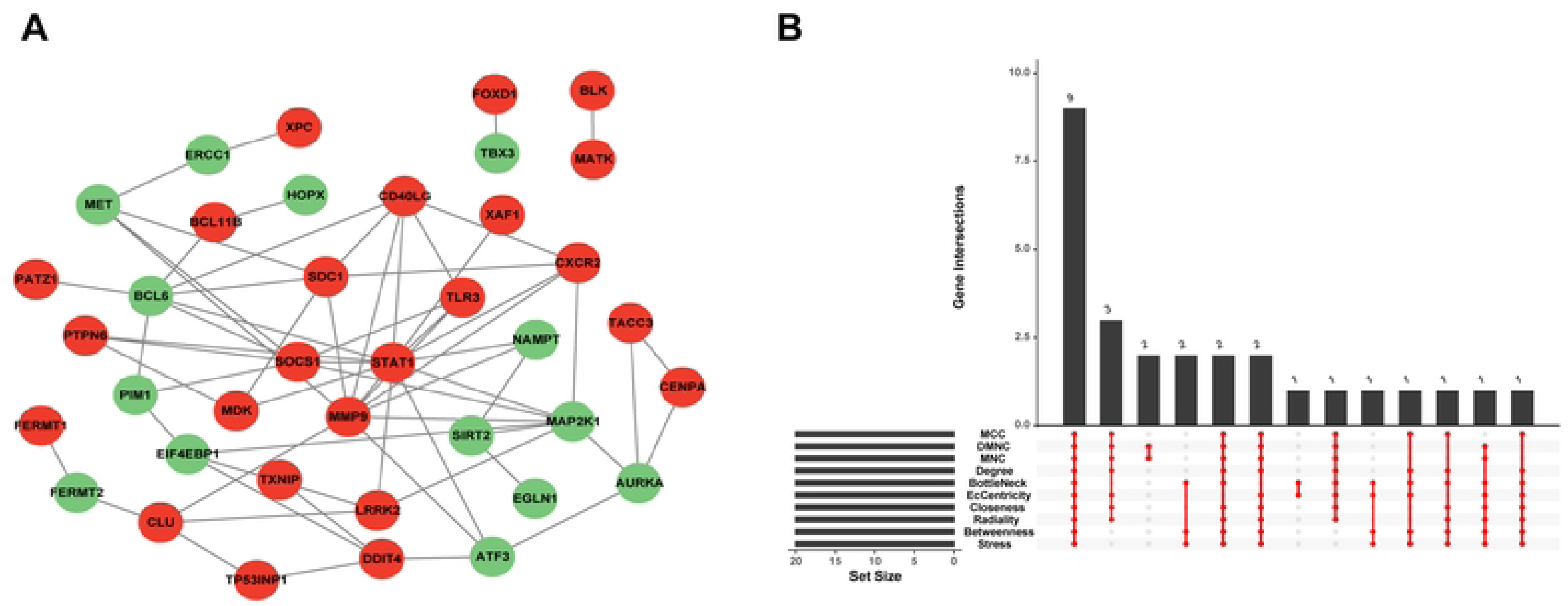
Protein–protein interaction (PPI) network analysis and identification of hub genes. **A** PPI network constructed from CS-DEGs. Red circles, upregulated DEGs; green circles, downregulated DEGs. **B** This graph shows the results of 12 algorithms used to screen out hub genes in CS-DEGs.

### Validation of the hub genes and correlation analysis

In the combined dataset comprising GSE84796 and GSE76701, all nine candidate hub genes demonstrated significant differential expression between CHF and normal samples. Specifically, *ATF3*, *EIF4EBP1*, *MAP2K1*, *MET*, and *NAMPT* exhibited higher expression in normal samples, whereas *MMP9*, *SDC1*, *SOCS1*, and *STAT1* were expressed at higher levels in CHF samples (Fig. 6A–I). The AUC exceeded 0.7 for all hub genes (Fig. 7A), indicating their significant diagnostic value for CHF. Coexpression analysis (Fig. 7B) revealed that *ATF3* and *NAMPT* exhibited the strongest positive correlation, with a correlation coefficient approaching 0.8 (Fig. 7C), whereas *EIF4EBP1* and *MET* displayed the strongest negative correlation (Fig. 7D).

**Fig. 6.**
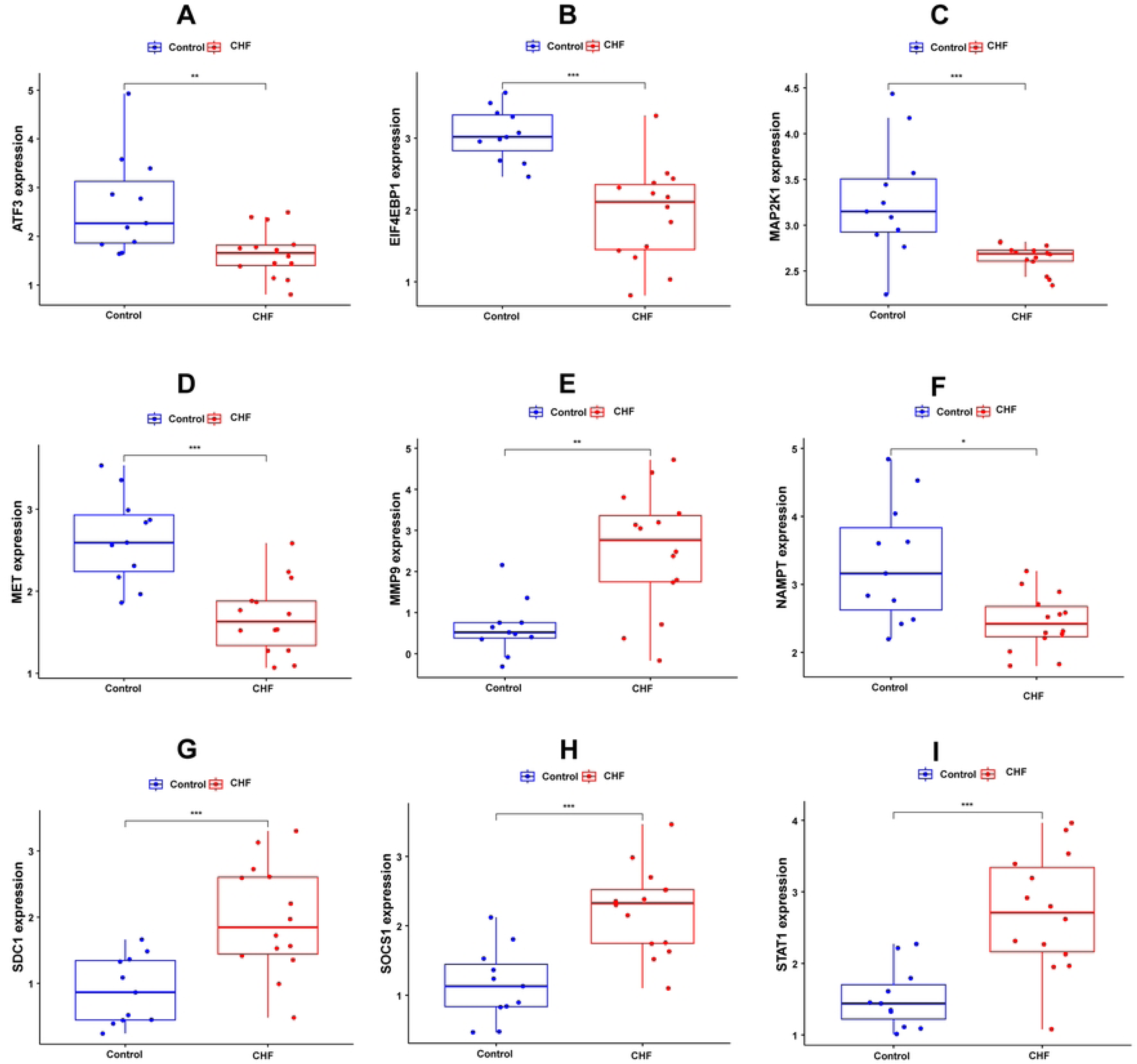
Expression profiles of nine hub genes in the combined microarray dataset (GSE76701 and GSE84796). The levels of expression were compared between the CHF and control groups. **A** *AFT3*, **B** *EIF4EBP1*, **C** *MAP2K1*, **D** *MET*, **E** *MMP9*, **F** *NAMPT*, **G** *SDC1*, **H** *SOCS1*, and **I** *STAT1*. \**P* < 0.05, \*\**P* < 0.01, \*\*\**P* < 0.001.

**Fig. 7.**
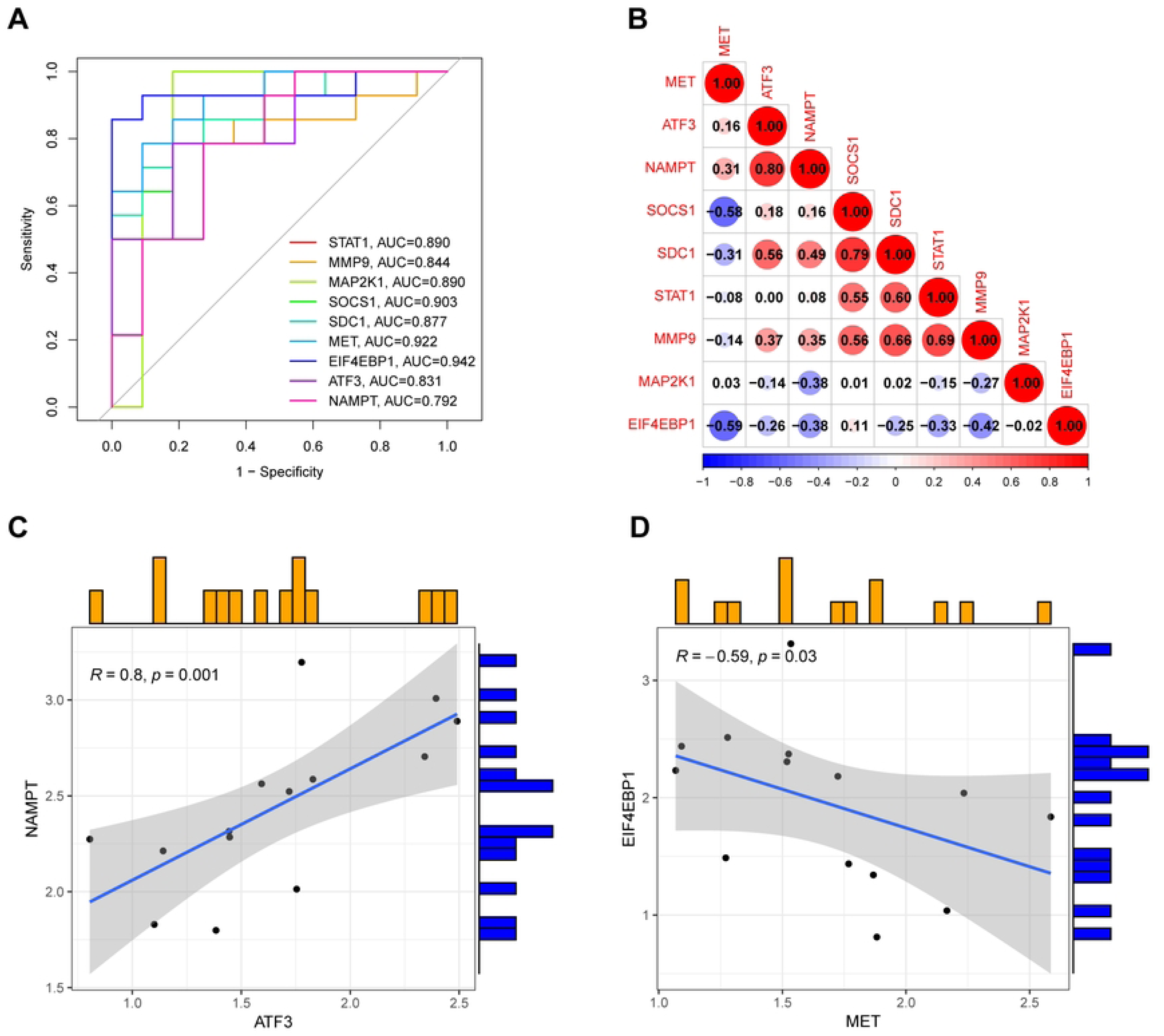
ROC curve and correlation analysis of hub genes. **A** The AUC values of the ROC curves for all hub genes exceed 0.7. **B** Correlation among the expression levels of various hub genes. **C** Correlation between *NAMPT* and *ATF3*. (D) Correlation between *EIF4EBP1* and *MET*.

### Construction of the hub gene–ceRNA–TF regulatory network

Utilizing the NetworkAnalyst database, we predicted the upstream regulatory miRNAs and lncRNAs, as depicted in Fig. 8A. Then, we predicted hub gene-related TFs. The interaction network comprised 7 hub genes and 11 TFs (Fig. 8B). Specifically, *MMP9* is regulated by eight TFs, including RELA and STAT3. *STAT1* is regulated by five TFs, including RELA and STAT3, whereas *SOCS1* is regulated by five TFs, such as STAT3 and STAT6. Additionally, NFKB1 and KLF6 target *ATF3*, and RELA and NFKB1 target *NAMPT*. Furthermore, TP53 and HIF1A were found to target *MET*, whereas RELA and NFKB1 target *SDC1*. Within the gene–TF network, RELA, NFKB1, and STAT3 exhibited close interactions with the hub genes.

**Fig. 8.**
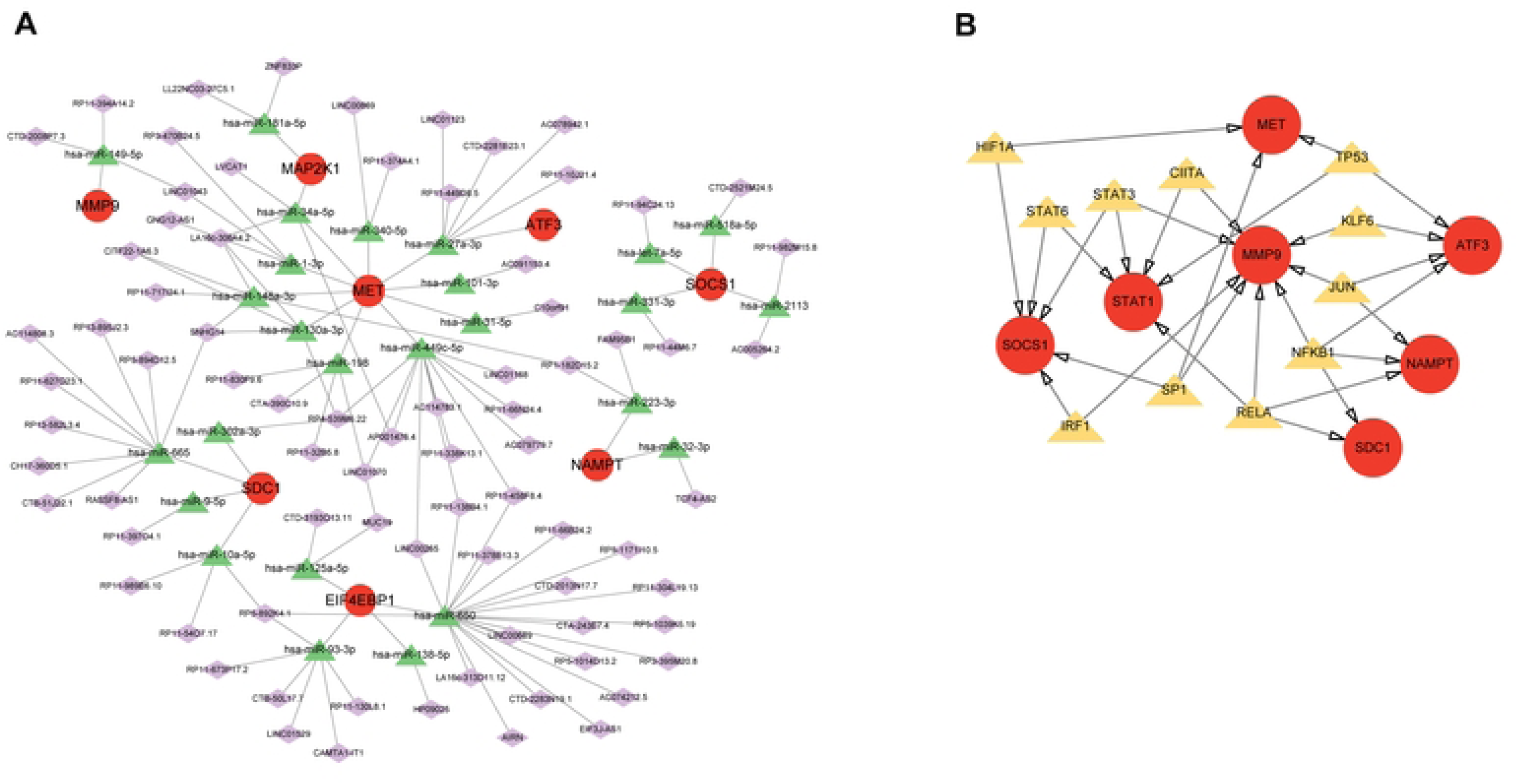
Construction of ceRNA network. **A** lncRNA–miRNA–mRNA network. Diamonds, lncRNA; triangles, miRNA; circles, mRNA. **B** Construction of hub gene– TF network. Circles, mRNA; triangles, TF

### Identification of potential therapeutic drugs for hub genes

We attempted to identify potential therapeutic drugs that regulate the expression of hub genes in CHF using DSigDB. In total, 42 candidate drugs for CHF treatment were identified (Supplementary Material S3). We identified drugs, including metformin and deguelin that target *MMP9*, 22 drugs, such as 2,4-dinitrofluorobenzene and indomethacin that target *ATF3*, 18 drugs, including rottlerin and bicalutamide that target *EIF4EBP1*, 20 drugs, namely rottlerin and bicalutamide, target *MAP2K1* expression, 13 drugs, such as tribenoside and butyl 4-hydroxybenzoate, target *MET*, 7 drugs, including 2,4-dinitrofluorobenzene and indomethacin, target *NAMPT*, 4 drugs, such as thalidomide and bortezomib, target *SDC1*, and 2 drugs (vanadium pentoxide and ciclopirox) target *SOCS1*. Figure 9A and 9B depicts the top 30 and top 10 relationship graphs between drugs and hub genes. However, we did not identify any potential drugs that targeted *STAT1* in this database. As the top-ranked drug, metformin was selected for molecular docking studies with the corresponding hub genes.

**Fig. 9.**
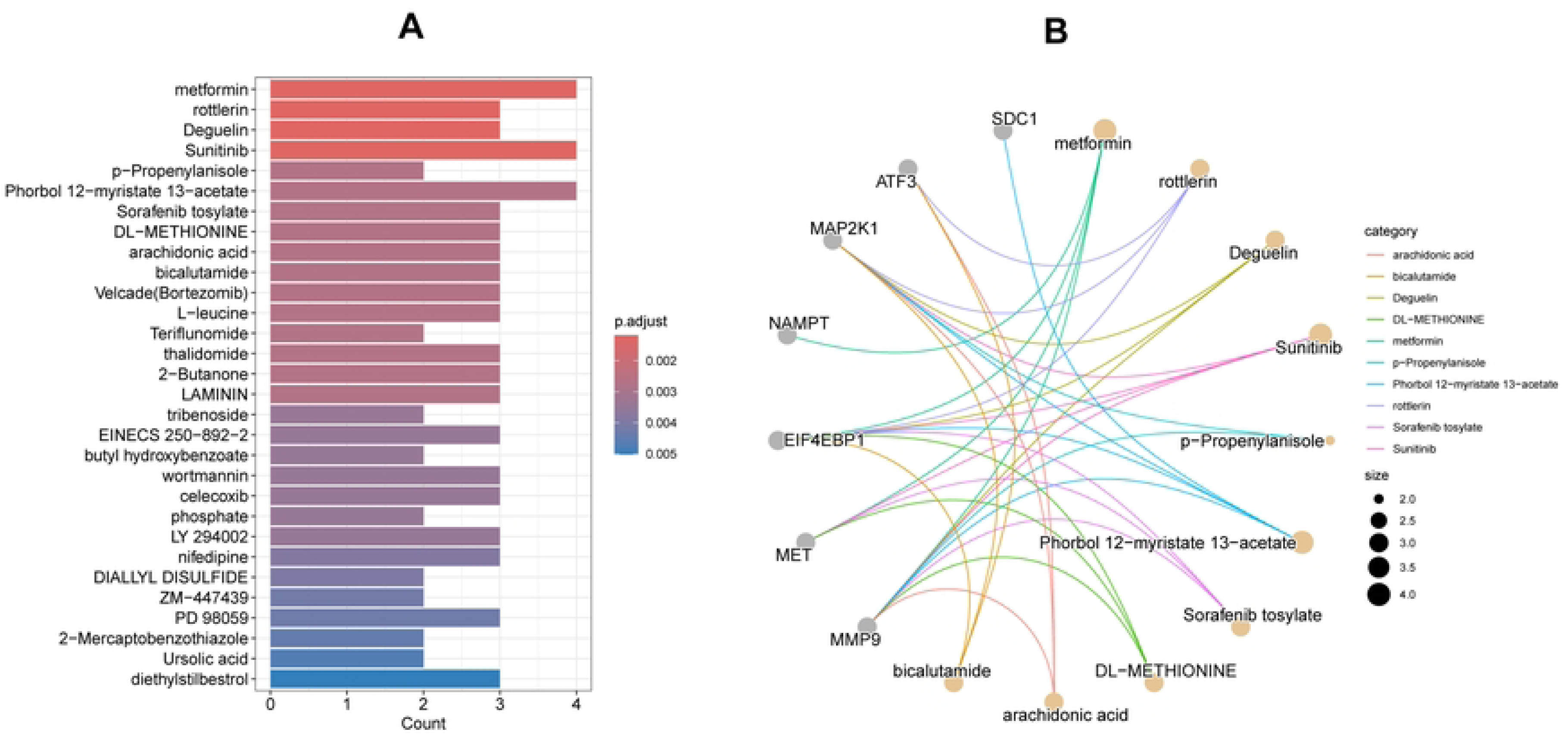
Prediction of the relationship between hub genes and drugs. **A** Top 30 drugs exhibiting the most significant enrichment. X-axis, drug names; y-axis, number of hub genes enriched by each drug. **B** Diagram showing the relationship between hub genes and drugs. Gray circles, hub genes; light yellow circles, drugs.

### Molecular docking of metformin with hub target proteins

We utilized molecular docking technology to investigate the interactions between metformin and four key target genes: *MMP9*, *MET*, *EIF4EBP1*, and *NAMPT*. Utilizing AutoDock software, our docking simulations yielded remarkably low binding energies (<−4.0 kcal/mol, Table 2), underscoring the potent binding affinity of these crucial proteins for metformin. Subsequently, we employed PyMOL to visualize the binding conformation with the lowest energy (Fig. 10). This visualization vividly illustrated the intimate relationship between metformin and its key targets, offering valuable insights into the structural basis of their interactions.

**Fig. 10.**
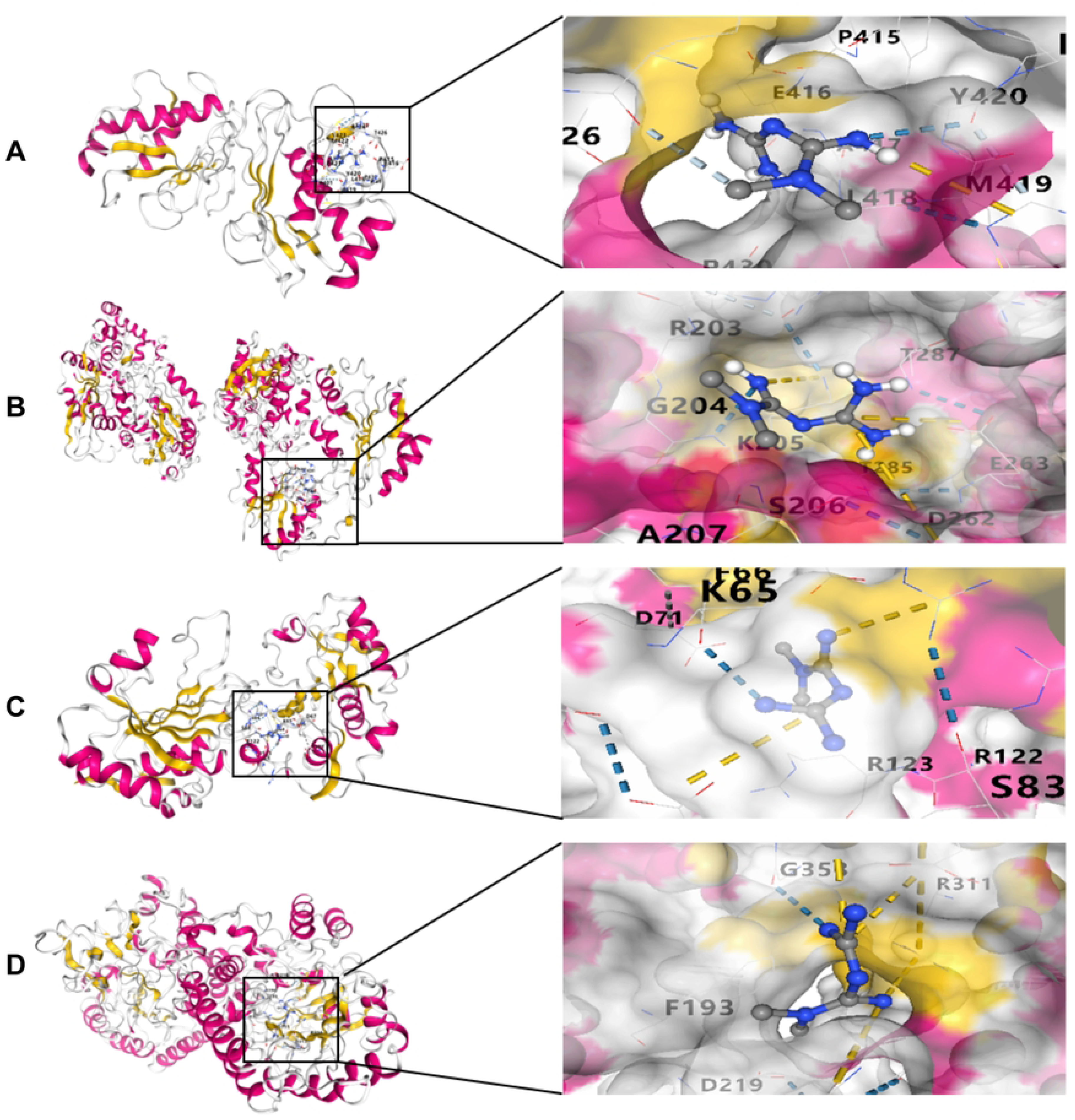
Molecular docking of target proteins with metformin. **A** Metformin and MMP9, **B** metformin and MET, **C** metformin and EIF4EBP1, and **D** metformin and NAMPT.

### Establishment of a rat CHF model

During the modeling period, the rats in the control group exhibited agile movement, a rapid response, glossy fur, and a good appetite. By contrast, the rats in the model group were typified by sluggishness, fatigue, dull fur, and a poor appetite. Body weight gradually increased over time in the control rats. Conversely, model rats exhibited slow weight gain in the first 2 weeks, followed by significant decreases in weight (*P* < 0.05, Fig. 11A). Compared with the findings in control rats, rats with CHF displayed significantly increased NT-pro BNP content in (Fig. 11B). The myocardial structure of the control group was neat and orderly, with regular cell morphology, uniform cell size, clear intercellular substance, no inflammatory cell infiltration, no necrosis, and no fibrous changes. The myocardial tissue was in a normal state (Fig. 11C). In the model group, the myocardial fiber structure was significantly disordered, displaying crisscrossing or breaks. Myocardial cells were swollen, with slight changes in the cell nucleus, widened intercellular substance, and visible infiltration of a few inflammatory cells, accompanied by local myocardial fibrotic changes. The myocardium was in a moderate injury state (Fig. 11D). These results support the successful establishment of the rat CHF model.

**Fig. 11.**
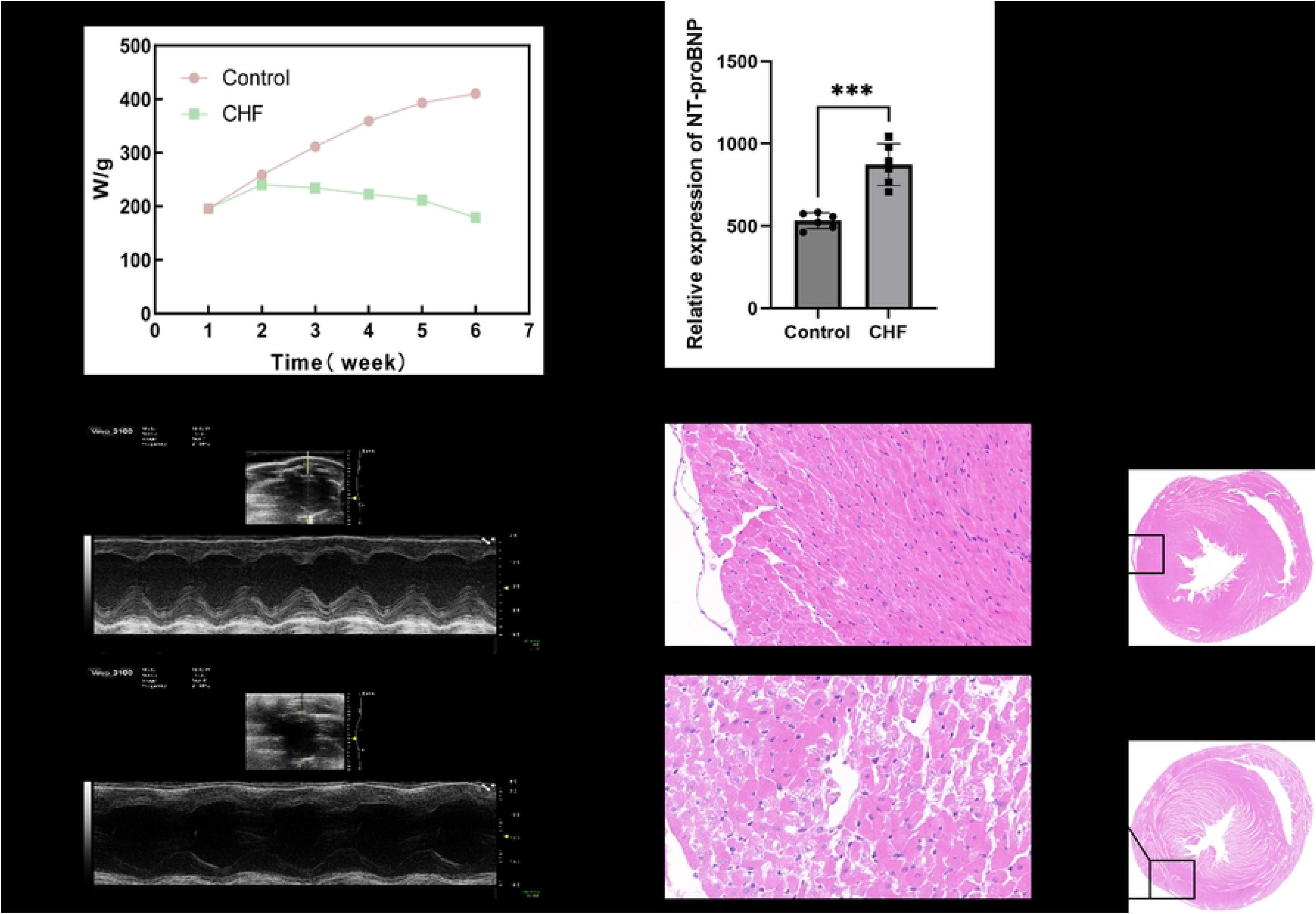
Results of experiments using a rat model of CHF successfully established via doxorubicin induction. **A** Weight change in rats in the CHF and control groups. **B** Comparison of NT-proBNP concentrations in blood samples collected from rats in the CHF and the control groups. **C** Comparison of ejection fractions between the CHF and control groups. **D** H&E staining of left ventricular tissues from the CHF and control groups. \*\*\**P* < 0.001.

### Validation of hub gene expression

Next, we conducted RT-qPCR to verify the expression of the hub genes. The primer sequences are listed in Table S3. Compared with that in the control group, *MMP9*, *SDC1*, *SOCS1*, and *STAT1* expression was significantly elevated in the CHF group, whereas *ATF3*, *EIF4EBP1*, *MAP2K1*, *MET*, and *NAMPT* were notably downregulated (Fig. 12A–I). *EIF4EBP1* and *MET* expression was inconsistent with that recorded in the previous database, possibly because of tissue heterogeneity. Nevertheless, the expression of the remaining hub genes aligned with that in the database.

**Fig. 12.**
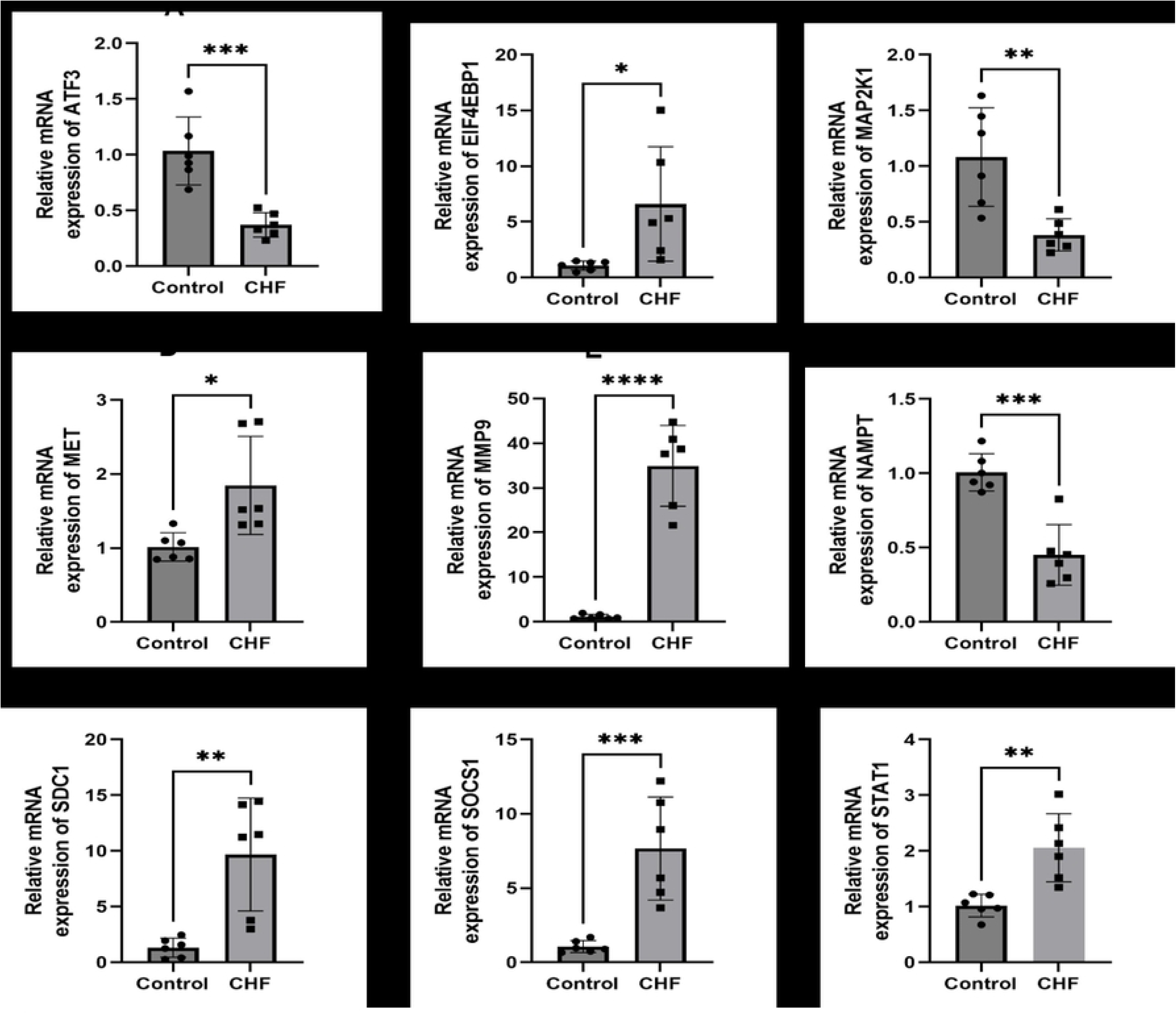
qRT-PCR assay determination of mRNA expression levels of nine hub CS-DEGs. **A–I** Comparison of the expression levels of *ATF3*, *EIF4EBP1*, *MAP2K1*, *MET*, *MMP9*, *NAMPT*, *SDC1*, *SOCS1*, and *STAT1* mRNA between the CHF and control groups. \**P* < 0.05, \*\**P* < 0.01, \*\*\**P* < 0.001, \*\*\*\**P* < 0.0001.

## Discussion

CHF is a complex clinical syndrome, and its complex, multidimensional pathogenesis involves multiple abnormalities in cardiac structure and function[31]. It is generally believed that the mechanisms underlying CHF mainly involve myocardial remodeling, abnormal myocardial energy metabolism, and excessive neuroendocrine activation. Current treatment methods mainly aim to reduce the cardiac load and regulate the neuroendocrine system. However, these methods lack long-term efficacy, and new treatment strategies are urgently needed[32]. As a self-defense mechanism, cellular senescence is closely related to various pathophysiological processes, playing an especially important role in the occurrence and development of heart failure. Its mechanism remains to be further explored[33, 34].

This study investigated the role of cellular senescence in CHF by combining transcriptome analysis and animal models to identify the characteristic genes associated with cellular senescence and explore potential therapeutic drugs. By analyzing public datasets (i.e., GEO, CellAge), we identified nine hub genes related to cellular senescence and established a good diagnostic model (with AUCs exceeding 0.7). In addition, we predicted the regulatory network of 94 lncRNAs and 63 miRNAs, which might play important roles in regulating the expression of hub genes and their biological functions. Our results provide new insights into the diagnosis and treatment of CHF, potentially opening up new avenues for improving patient outcomes.

This study identified nine hub genes associated with cellular senescence with significant roles in the pathogenesis of CHF. The changes in the expression of genes such as *STAT1*, *MMP9*, *MAP2K1*, *SOCS1*, *SDC1*, *MET*, *EIF4EBP1*, *ATF3*, and *NAMPT* might affect the structure and function of the heart, thereby influencing CHF progression. Previous studies reported that the TF STAT1 plays a crucial role in the stress response of the heart, and its activation is closely associated with cardiac dysfunction[35]. *MMP9* is involved the process of myocardial remodeling, and its upregulation in heart failure might promote changes in the structure of the heart and deterioration of its function[31, 36]. Additionally, *MET* plays an important role in cardiac regeneration and repair, and changes in its expression might be closely related to the aging process of the heart[37]. By analyzing the interactions and signaling pathways of these genes, we might obtain a deeper understanding of the pathogenesis of CHF and elucidate new targets for future treatment strategies.

We predicted 42 potential therapeutic drugs, among which metformin exhibits a significant binding effect with hub genes, providing new insights for the pharmacological treatment of CHF. Existing evidence indicates that metformin has a protective effect in multiple cardiovascular diseases, influencing cardiac function by improving the metabolic status and inhibiting inflammatory responses[31, 38]. Meanwhile, further exploration of the mechanisms of action of other candidate drugs is required to assess their potential application in the treatment of CHF[33]. Therefore, future research should evaluate the safety and effectiveness of these drugs in clinical practice to ensure that they can provide tangible therapeutic benefits to patients with CHF.

By establishing regulatory networks of miRNAs, lncRNAs, and TFs for hub genes, we have provided a new perspective for understanding the regulatory mechanisms of these genes in CHF. These noncoding RNAs might have pivotal roles in regulating the expression and biological functions of hub genes[39]. Notably, the regulatory capability of miRNAs can influence the transcription levels of hub genes, thereby further affecting CHF progression[40, 41]. Future research should explore the utility of modulating these noncoding RNAs in improving the pathological state of CHF, thus offering novel therapeutic insights. Verifying the biological functions of these networks will be a crucial direction for future research, aiming to provide more precise strategies for the treatment of CHF.

Experimental verification in a rat CHF model revealed that the expression of some hub genes did not align with expectations, suggesting potential differences in gene expression regulation across different experimental models. For instance, the expression of *EIF4EBP1* and *MET* did not align with the database results, potentially reflecting limitations in the experimental design or model selection. Therefore, improving the experimental design to ensure that the model better simulates the pathological state of CHF in humans will be a key focus of future research. Comprehensively considering these factors will improve our understanding of the regulatory mechanisms of hub genes in CHF, thereby providing a more reliable basis for clinical practice.

The limitations of this study are primarily reflected in the relatively small sample size and the lack of broader clinical validation. Although we identified hub genes related to cellular senescence by analyzing the dataset, the insufficient sample size might have led to insufficient statistical significance of the results. Additionally, interbatch differences in the dataset might have also affected the reliability of the results. Although we proposed potential therapeutic drugs in the study, the lack of wet experiment validation requires further exploration of the biological significance and clinical applicability of the results. Therefore, future research should expand the sample size and combine multicenter clinical data to enhance the universality and reliability of the results.

## Conclusions

In summary, this study successfully identified nine hub genes and potential therapeutic drugs through a comprehensive analysis of cell senescence-related genes in patients with CHF. This discovery provides a new perspective for the diagnosis and treatment of heart failure and highlights directions for future research. Further validation and intensive study of these hub genes and their regulatory networks could improve patient prognosis by promoting the development of clinical management and treatment strategies for CHF.

## List of abbreviations

AUC: Area under the curve
BP: Biological process
CC: Cellular component
CHF: Chronic heart failure
DEGs: Differentially expressed genes
CS-DEGs: cell senescence-related DEGs
GEO: Gene Expression Omnibus
GO: Gene Ontology
KEGG: Kyoto Encyclopedia of Genes and Genomes
MF: Molecular function
ROC: Receiver operating curve
TF: Transcription factor

## Acknowledgements

Not applicable.

## Authors’ contributions

Hao Xie and Zuyuan You wrote and revised the paper. Qi Yang, Juyi Wan, and Mingbin Deng investigated information. Bin Liao, Mingbin Deng and Qi Yang designed the study.

## Data Availability

On request, the corresponding author will provide access to the data used to support the study’s conclusions.

## Ethical approval

This study was approved by the Ethics Committee for Animal Experiments of Southwest Medical University.The Ethical approval number is 20250530-001.

